# Obesity reprograms adipose extracellular vesicles to induce muscle atrophy via miR-150-5p-mediated transcriptional silencing

**DOI:** 10.64898/2025.12.09.693129

**Authors:** Joshua MJ Price, Michael Macleod, Thomas Nicholson, Caitlin M Ditchfield, Bethy Airstone, Natalie Lachlan-Jiraskova, Edward T Davis, Kostas Tsintzas, Simon W Jones

## Abstract

**Background:** Sarcopenic obesity, where excess body fat coexists with reduced muscle mass and function, is becoming increasingly common in ageing populations and contributes to poor physical and metabolic health. Although adipose tissue-secreted factors are implicated in muscle decline, the specific mechanisms remain unclear. Extracellular vesicles (EVs), which carry regulatory cargo such as microRNAs (miRNAs) between cells, may play a key role in this adipose-muscle communication.

**Methods:** EVs were isolated from adipose-conditioned media (ACM) collected from lean and non-lean human donors using ultracentrifugation. Donors were grouped by BMI (lean: 20.7–24.4; non-lean: 25.3–39.3) and age (younger: 31–56 years; older: 60–84 years). EVs were characterised using nanoparticle tracking analysis (NTA), ExoView, nanoscale flow cytometry (CytoFLEX Nano), and transmission electron microscopy (TEM). Primary human myoblasts were differentiated into myotubes and treated for 24 hours with lean or non-lean EVs (1.3×10⁹ particles/ml) or left untreated. Myotube thickness was measured by immunofluorescence microscopy. Transcriptomic changes were assessed by bulk RNA sequencing. EV miRNA cargo was profiled by small RNA-seq and validated by qPCR. The role of miR-150-5p was tested using antagomir inhibition.

**Results:** Non-lean EVs significantly reduced myotube thickness compared to both untreated controls (8.7 ± 1.66 µm vs. 12.4 ± 1.72 µm, p < 0.01) and lean EV-treated myotubes (8.7 ± 1.66 µm vs. 13.2 ± 3.84 µm, p < 0.05), indicating a donor BMI-specific effect. This atrophy was restricted to myotubes derived from older donors. MAFbx expression was significantly increased in response to non-lean EVs (p < 0.05). RNA-seq revealed 471 differentially expressed genes (DEGs) in EV-treated vs. untreated cells and 293 DEGs between lean and non-lean EV conditions, with enrichment in inflammatory (TNF, IL1B), oxidative stress, mitochondrial, and chromatin pathways. Small RNA-seq identified 7 differentially expressed miRNAs, including miR-150-5p and miR-193b-5p, both significantly upregulated in non-lean EVs and validated by qPCR. Inhibiting miR-150-5p partially rescued myotube thickness (10.5 ± 1.37 µm vs. 8.7 ± 1.66 µm, p < 0.05) and reduced MAFbx expression.

**Conclusions:** EVs from non-lean adipose tissue drive muscle atrophy and transcriptional changes in an age-dependent manner. These effects are partially mediated by miR-150-5p, highlighting a mechanistic role for EV cargo in adipose-muscle signalling. Targeting EV-derived miRNAs may offer a novel strategy to combat muscle loss in obesity and ageing.

## 1. Introduction

Adipose tissue is a complex endocrine organ, comprising a heterogeneous population of cells including adipocytes, immune cells, fibroblasts, and endothelial cells [1]. As a major source of adipokines such as leptin and resistin, the cellular composition and phenotype of adipose tissue exerts influence over the pathology of distal tissues by modulating metabolic and inflammatory processes [2]. Excess adiposity is associated with insulin resistance, systemic inflammation and multimorbidity; effects that are compounded by age-related loss of skeletal muscle mass and function (sarcopenia) and contribute to progressive decline in metabolic health [2–4]. The comorbidity of obesity and sarcopenia (termed sarcopenic obesity) is becoming increasingly prevalent, and severely compromises mobility, physical independence, and quality of life [5]. Investigating the nature of this interplay is challenging due to confounding covariates, including age, physical activity levels, sex and multimorbidity [6]. Common inflammatory comorbidities such as chronic liver disease [7], chronic kidney disease [8] and arthritis [9] exacerbate the sarcopenic phenotype.

Obesity further disrupts homeostatic signalling, creating a pro-atrophic environment that may accelerate sarcopenia [6]. Pro-inflammatory cytokines and adipokines such as TNFα and resistin activate muscle atrophic signalling pathways [10] via the E3 ubiquitin ligases Atrogin-1 (MAFbx) and MuRF1, which promote proteasomal degradation of muscle proteins and has been reviewed extensively [11].

We have previously reported that the secretome of adipose tissue from non-lean individuals impairs myogenesis of older adult human skeletal muscle, resulting in muscle fibres of reduced thickness [10]. Adipose secretome-induced impairment in myogenesis was not observed in muscle from young adults [10]. This suggests that with age and obesity, adipose tissue secretome drives pathological cross-talk with skeletal muscle. However, the precise intracellular and intercellular mechanisms by which excess adipose tissue promotes skeletal muscle ageing remain poorly understood.

Extracellular vesicles (EVs) are small, phospholipid bound particles that cannot replicate. EVs contain diverse biological cargo including proteins, lipids and nucleic acids [12]. They are emerging as pivotal mediators of intercellular communication, particularly through the transfer of miRNAs; small, non-coding RNAs that regulate gene expression post-transcriptionally [13]. Adipose tissue is a major source of both EV [14] and EV-associated miRNA [15, 16], with evidence of communication to both the liver and to skeletal muscle [16]. Indeed, miR-155 is found in adipose from non-lean individuals and promotes inflammation through inhibition of C/EBPβ (CCAAT/enhancer-binding protein beta) and induction of NF-κB via targeting of SOCS1 [17]. miR-155 knockout mice exhibit protection against obesity-induced insulin resistance and glucose intolerance under a high-fat diet [18].

While the total concentration of circulating EVs increases with BMI, direct attribution to adipose tissue or specific changes in adipose biology has been challenging as many studies focus on isolated individual cellular populations from adipose tissue, potentially missing the collective contribution of its diverse cell types [19]. However, evidence shows that the concentration of EVs released into the adipocyte secretome is positively correlated with donor adiposity, with obese individuals exhibiting higher levels of adipocyte-derived EVs compared to lean individuals [19].

Given the critical role of EVs in mediating intercellular communication, and evidence of adipose-muscle crosstalk in driving inflammatory phenotypes [20], we investigated the role of non-lean ACM-derived EVs in promoting muscular atrophy. We aimed to profile human ACM-derived EVs, including their microRNA cargo and characterise their role in driving muscle atrophy to identify novel mechanisms of atrophy.

## 2. Methods

### 2.1 Skeletal muscle and adipose tissue collection

Skeletal muscle (∼ 200 mg) and adipose tissue (∼ 2000 mg) samples were collected peri-operatively from adults undergoing elective orthopaedic surgery at either The Royal Orthopaedic Hospital (Birmingham, UK) or Russell’s Hall Hospital (Dudley, UK). Ethical approval was provided by the UK National Research Ethics (16/SS/0172). Donors were categorised by BMI, as lean (BMI 18.5–24.9; observed range: 20.7–24.4) or non-lean (BMI ≥25; observed range: 25.3–39.3).For age, donors were grouped as younger (<60 years; 31–56 years) or older ((≥60 years; 60–84 years). This stratification aligns with literature distinguishing older and younger adults [21].

### 2.2 Quantification of myotube thickness

Details on primary human myoblast isolation and differentiation are provided in Supplementary Methods. Myotube thickness quantification was conducted on myotubes cultured and differentiated in 24-well plates. The culture medium was removed, and myotubes were fixed with 2% (para)formaldehyde in PBS for 30 min at room temperature (RT). Cells were then permeabilised with 100% methanol for 10 min, followed by blocking with 5% goat serum (Gibco™, Thermo Fisher, #16210064) in PBS for 30 min at RT.

Subsequently, myotubes were incubated with the primary antibody (Rabbit polyclonal anti-desmin antibody, Abcam, #ab15200, 1:500 in 1% BSA/PBS) overnight at 4°C, followed by incubation with the secondary antibody (Goat anti-Rabbit IgG (H + L) Cross-Absorbed, FITC-conjugated, Thermo Fisher, #F-2765, 1:800 in PBS) for 1 hour in the dark at RT. Wells were then washed with PBS, after which DAPI staining solution (1:5000 in PBS, Abcam, #ab228549) was added for 5 min in the dark. Finally, wells were washed again with PBS, before Fluoromount Aqueous Mounting Medium (Sigma, Cat# F4680)and coverslip were applied. Immunofluorescent-stained myotubes were imaged using an epifluorescence/brightfield microscope (Leica DMI6000,10 images per well were captured with a ×63 objective). Image analysis was performed using ImageJ software, with MTT calculated as the average of five cell width measurements taken along the length of each myotube.

### 2.3 Generation of adipose conditioned media and EV isolation

Subcutaneous adipose tissue (SAT) was gently washed with PBS, cut with a scalpel into 2-3 mm^2^ pieces and placed in a 50ml falcon tube and incubated in DMEM at a ratio of 1 g tissue to 10 mL medium for 24 hours at 37 °C, 21% O_2_ and 5% CO_2_. After 24 hours the adipose conditioned medium (ACM) was removed, filtered through a 0.22uM filter (Millipore) and double-centrifuged at 2000 x *g* for 20 min. ACM was stored at −80°C before being thawed for use. EV were isolated from ACM by ultracentrifugation at 100,000 x *g* for 16 hours at 4°C. The resultant pellet was resuspended in 0.22uM filtered PBS at 10x concentrate (i.e. EV derived from 4ml of ACM were resuspended in 400ul of PBS).

### 2.4 EV Characterisation

EVs were diluted 1:100 in DPBS (Thermo Fisher Scientific, Cat# 14190094) and characterised by nanoparticle tracking analysis (NTA) using the NS Pro (Malvern Panalytical, Malvern). Settings were automated using the NSExplorer software. Tetraspanin characterisation of EVs was also performed using the ExoView R100 (Unchained Labs) as previously described [22]. Samples were diluted at 1:100 in Incubation Solution (Unchained Labs).

EV size and concentration were also measured using the CytoFLEX Nano (Beckman Coulter), and morphology confirmed by transmission electron microscopy (TEM) using a JEOL 1400Flash with Gatan OneView camera (100 kV). Detailed protocols are available in the Supplementary Methods. All relevant information has been shared with EV TRACK ID: EV250060 [23].

### 2.5 RNA isolation, RNA sequencing analysis and PCR

Primary myotubes were treated with PBS vehicle control (n = 5) or EV isolated from adipose conditioned media (n = 3 lean, n = 8 non-lean). Total RNA was isolated using TRIzol reagent (Life Technologies, UK). The mean average RNA integrity number (RIN) of myotube RNA was 9.2 (SD = 1.2, Agilent Bioanalyzer). Library preparation and bulk RNA sequencing were performed by Lexogen NGS Services (Vienna, Austria). Subsequent analysis used Bowtie2 against the hg38 reference genome. DESeq2 identified differentially expressed transcripts (p value <0.05, fold change >1.5 or <-1.5). RNA from ACM-derived EV was isolated with mRNeasy Tissue/Cells Advanced Micro Kit (Qiagen, UK) with on-column DNA digestion following manufacturer guidelines. Library preparation and small RNA sequencing were performed by BGI (Shenzhen, China). Subsequent analysis was undertaken using MiRDeep2 against miRNA precursors and mature sequences using miRBase as a reference (PMID: 30423142). DESeq2 was used to identify differentially expressed transcripts (p value <0.05, fold change >1.5 or <-1.5).

Validation by qPCR is described in Supplementary Methods.

### 2.6 Extracellular Vesicle and Antagomir Treatment

Myotubes were treated with ACM-derived EV at a final concentration of 1.3 × 10^9^ particles/mL for 24 hours. A PBS vehicle control was included as the untreated condition. In some experiments, ACM or the EV-depleted supernatant following EV isolation was used instead of EVs. For all ACM-based treatments, wells received 50% ACM and 50% differentiation media.

In separate experiments, myotubes were treated with EVs derived from non-lean donors in the presence and absence of an antagomir against miR-150-5p (#HYRI00301A, MedChemExpress). The antagomir, modified with a 3’ cholesterol tag, was administered simultaneously with EVs at a final concentration of 0.5 μM.

### 2.7 Statistical Analysis

All statistical analyses and figure generation were performed in RStudio (R version 4.3.1). Data distribution was assessed visually using histograms and formally using the Shapiro-Wilk test, and parametric or non-parametric tests were used as appropriate. For comparisons between two groups, either Student’s *t*-test or the Wilcoxon rank-sum test was used. For multiple group comparisons, one-way ANOVA was followed by pairwise *t*-tests with Holm-Bonferroni correction, or Kruskal-Wallis test followed by Wilcoxon rank-sum tests with Holm-Bonferroni correction. Statistical significance was defined as *p* < 0.05.

## 3. Results

### 3.1 The characterisation of EVs released from lean and non-lean human adipose tissue

The successful isolation of EVs from ACM by ultracentrifugation was confirmed through a multi-modal characterization approach, including nanoparticle tracking analysis (NTA), ExoView, nanoscale flow cytometry, and transmission electron microscopy (Figure 1A, Supplementary Figure 1). TEM images (Supplementary Figure 1B) show vesicles with spherical structures enclosed by a lipid bilayer membrane, consistent with the expected morphology of EVs. No morphological differences were observed between lean and non-lean samples.

**Figure 1.**
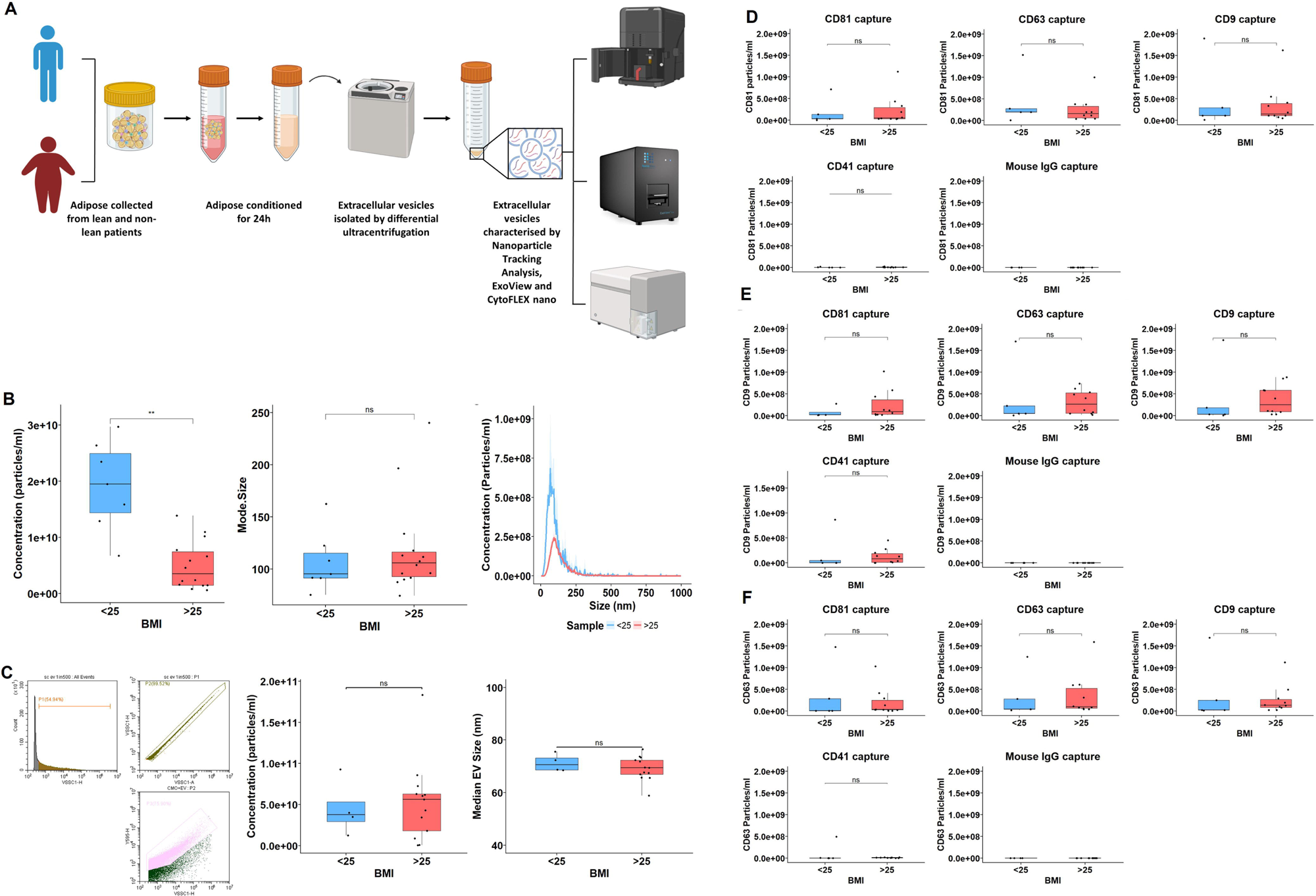
Quantitative and phenotypic characterization of ACM-derived extracellular vesicles (EVs) isolated from lean and non-lean conditioned media. **(A)** Schematic workflow for EV isolation and characterisation. Adipose tissue was digested to generate adipose-conditioned media (ACM), from which extracellular vesicles (EVs) were isolated by differential centrifugation and analysed using nanoparticle tracking analysis (NTA), CytoFLEX Nano flow cytometry, and ExoView. **(B)** NTA revealed significantly lower EV concentrations in non-lean ACM (BMI >25, n = 14) compared to lean ACM (BMI <25, n = 7; unpaired t-test). No significant differences were observed in mode EV size (Wilcoxon test). Representative size distribution curves show the average NTA trace across all lean (n = 7) and non-lean (n = 14) samples. **(C)** Gating strategy for EV detection by CytoFLEX Nano using violet side scatter (VSSC1). A small-scale trial using CellMask Orange (CMO) confirmed that the majority of detected particles were lipid-positive. Quantitative analysis across the full cohort (lean n = 4, non-lean n = 13) revealed no significant differences in EV concentration or size (Wilcoxon tests). **(D–F)** ExoView analysis of ACM-derived EVs captured on antibody-coated chips and detected via fluorescent tetraspanin markers: CD81–AF555 (**D**), CD63–AF647 (**E**), and CD9–AF488 (**F**). EVs were captured using anti-CD81, CD63, CD9, CD41, or mouse IgG control spots. No significant differences in tetraspanin-positive EV counts were observed between lean (n = 5) and non-lean (n = 10) samples (Wilcoxon tests). All boxplots display the median and interquartile range (IQR); whiskers extend to 1.5×IQR, and individual donor values are shown.

NTA revealed that EVs from non-lean ACM (5.04 × 10⁹ ± 4.30 × 10⁹ particles/ml) were significantly reduced in concentration compared to those from lean ACM (1.92 × 10¹⁰ ± 8.02 × 10⁹ particles/ml, p < 0.01; Figure 1B). However, modal EV size did not differ significantly between groups (95.2 ± 23.8 nm vs. 105.8 ± 23.85 nm; p > 0.05), indicating a similar size distribution profile (Figure 1C).

ExoView analysis using tetraspanin-targeting antibodies (CD9, CD63, CD81) confirmed the presence of classical EV markers in both lean and non-lean samples, with no detectable differences in capture or fluorescence signal patterns across conditions (Figure 1D–F). Nanoscale flow cytometry similarly showed no group differences in EV size aligning more closely with ExoView than with NTA.

Together, these findings confirm successful EV isolation from ACM and show that while total EV concentration may be reduced from non-lean donors (as indicated by NTA), vesicle size and marker expression remain consistent across BMI groups.

### 3.2 Non-lean adipose EVs drive atrophy of human myotubes

Having confirmed the isolation of EVs from both non-lean and lean ACM, we then sought to determine their effect on human muscle. For these cross-talk studies, EV concentrations, as determined by NTA, were used to inform dilutions required for the treatment of myotubes. The chosen EV concentration (1.3×10^9^ particles/ml) for treatments was based on physiological relevance and sample size availability [12].

Myoblasts were differentiated to multi-nucleated myotubes over 8 days. Differentiated myotubes from young (Range = 31-56 years old) or old (Range = 60-84 years old) were treated for 24 hours with either control media, whole ACM (lean or non-lean), isolated non-lean EVs, or EV-depleted ACM (Figure 2A). In myotubes derived from young donors, TNF-α treatment trended toward reduced thickness, but no differences were observed across the ACM or EV conditions (Figure 2B).

**Figure 2.**
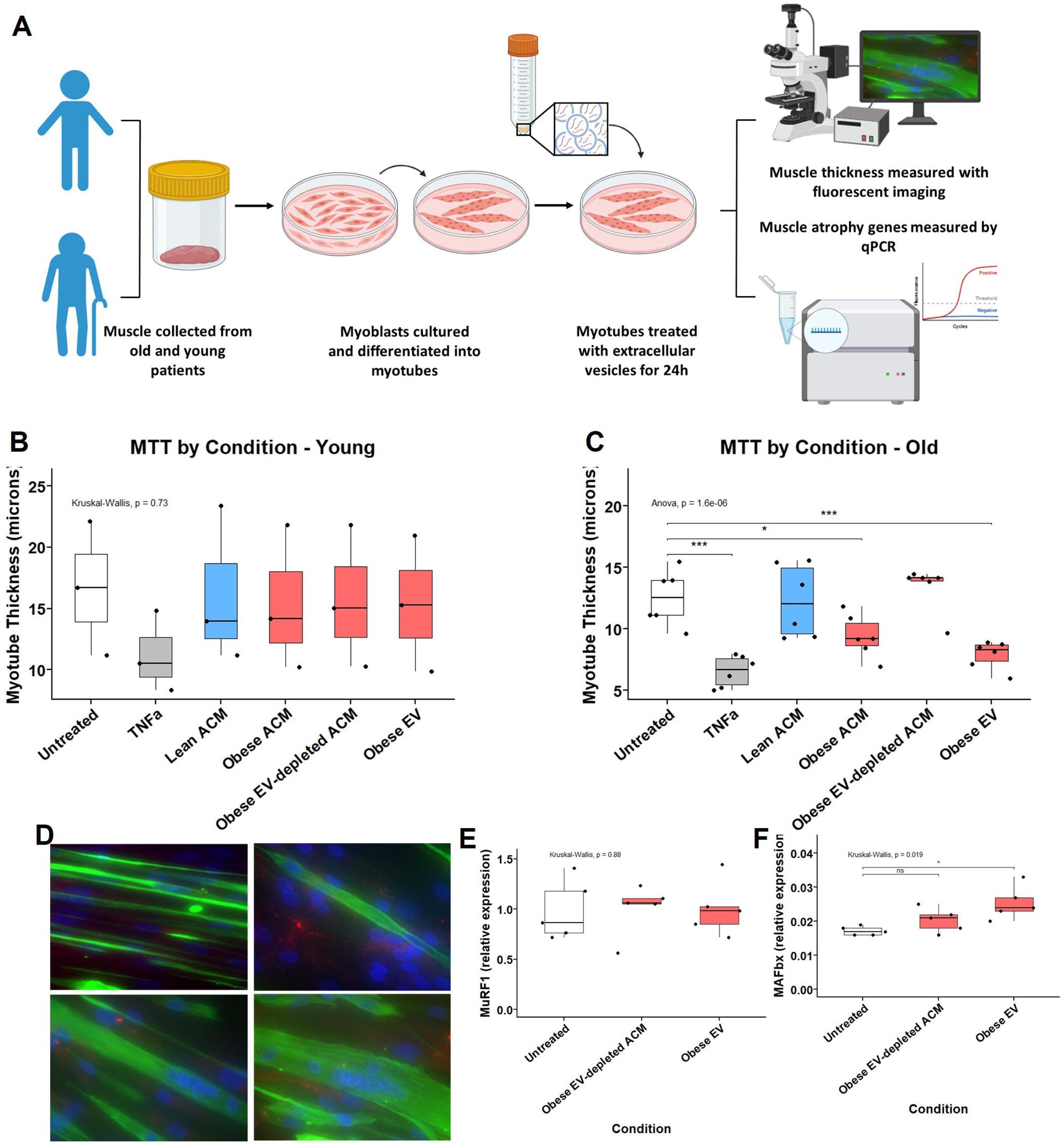
Non-lean ACM-derived extracellular vesicles (EVs) induce myotube atrophy in aged human muscle. **(A)** Schematic of experimental workflow. Human myoblasts derived from young and old donors were differentiated into multinucleated myotubes over 8 days, then treated for 24 hours with: untreated growth media, TNF-α (positive control), lean ACM, non-lean ACM, EV-depleted non-lean ACM, or isolated non-lean EVs. Myotube thickness was quantified by fluorescent imaging, and atrophy gene expression assessed by qPCR. **(B)** Myotube thickness in young donor-derived myotubes showed no significant differences across conditions (Kruskal–Wallis, p = 0.73). n = 3 young donors; all conditions were tested in cells from each donor. TNF-α-treated myotubes showed a consistent, though non-significant, reduction in thickness relative to untreated. **(C)** In old donor-derived myotubes (n = 6 donors), one-way ANOVA revealed a significant overall effect of treatment (p = 1.6e–06). Holm–Bonferroni corrected post-hoc comparisons showed that TNF-α significantly reduced myotube thickness compared to all other groups (***p < 0.001). Non-lean ACM also reduced thickness compared to untreated (*p < 0.05), while lean ACM had no effect. EV-depleted ACM failed to reproduce the atrophic effect, whereas isolated non-lean EVs significantly reduced myotube thickness compared to untreated (**p < 0.01). **(D)** Representative immunofluorescence images of desmin-positive myotubes (green) and nuclei (blue) across treatment conditions. **(E–F)** Expression of muscle atrophy genes in old donor-derived myotubes. **(E)** *MuRF1* expression did not significantly differ between groups (Kruskal–Wallis, p = 0.88). **(F)** *MAFbx* expression was significantly upregulated in response to non-lean EV treatment (Kruskal–Wallis, p = 0.019). Holm–Bonferroni corrected post-hoc Wilcoxon test confirmed a significant increase in the non-lean EV group compared to untreated controls (*p < 0.05), but not in the EV-depleted ACM condition. n = 5 donors per group; all conditions were tested in cells from each donor. qPCR values represent the average of technical triplicates per donor and were normalised to housekeeping genes (see Methods). All boxplots display the median and interquartile range (IQR); whiskers extend to 1.5×IQR, and individual donor values are shown.

In contrast, in myotubes derived from old donors, there was a significant (ANOVA, p < 0.0001) overall treatment effect (Figure 2C). Similarly to previous work on murine myotubes [24], we found TNF-α to be atrophic to human myotubes, with a significant reduction in myotube thickness observed in TNF-α treated myotubes, compared to untreated (5.85 ± 1.72 μm vs. untreated, p < 0.0001) Treatment with non-lean ACM significantly reduced myotube thickness compared to untreated controls (10.1 ± 1.07 μm vs. 12.4 ± 1.72 μm, p < 0.05), representing a ∼19% reduction, whilst lean ACM had no significant effect (13.2 ± 3.84 μm vs. 12.4 ± 1.72 μm, p > 0.05). EV-depleted non-lean ACM was also not significantly different from the untreated condition (12.0 ± 2.89 μm vs. 12.4 ± 1.72 μm, p > 0.05), whereas the isolated EV fraction from non-lean ACM induced a ∼30% reduction in myotube thickness (8.7 ± 1.66 μm vs. 12.4 ± 1.72 μm, p < 0.01), indicating that EVs are the active component mediating the atrophic effect. Representative desmin-stained images highlighting the differences in myotube thickness are shown in Figure 2D.

With the indication that the EVs from non-lean adipose tissue were the active component in driving atrophy, we next examined whether they modulated the mRNA expression of the muscle-specific ubiquitin ligases (atrogenes) MAFbx (Atrogin-1) and MuRF1. Expression of MuRF1 (Figure 2E) was not changed in myotubes treated with either non-lean EVs compared to control (Figure 2E). However, the expression of MAFbx was significantly upregulated in response to non-lean EVs (p <0.05), but not in response to EV-depleted supernatant (Figure 2F), providing further evidence of the muscle atrophic-mediating effect of non-lean adipose-derived EVs in older muscle.

### 3.3 EVs from non-lean adipose differentially modulate the transcriptome of human myotubes

To further assess how ACM-derived EVs influence skeletal muscle gene expression, we treated differentiated human myotubes (Donor age = 70 year old) to EVs from lean or non-lean adipose tissue for 24 hours and subjected them to bulk RNA sequencing. Transcriptomic profiling revealed widespread gene expression changes in response to EV treatment, with distinct signatures depending on donor adiposity (Figure 3A). A total of 471 DEGs (log₂FC ≥ 0.58 and p < 0.05) were identified between EV-treated and untreated myotubes, including 311 upregulated and 156 downregulated genes (Figure 3B). In the comparison between non-lean and lean EV treatments, 293 DEGs were identified, comprising 257 upregulated and 41 downregulated genes (Figure 3B). Principal component analysis (PCA) showed distinct clustering of untreated, lean EV-treated, and non-lean EV-treated samples, confirming that both EV exposure and patient characteristics influence transcriptional changes (Figure 3C). Heatmap analysis further demonstrated treatment-specific expression patterns (Figure 3D).

**Figure 3.**
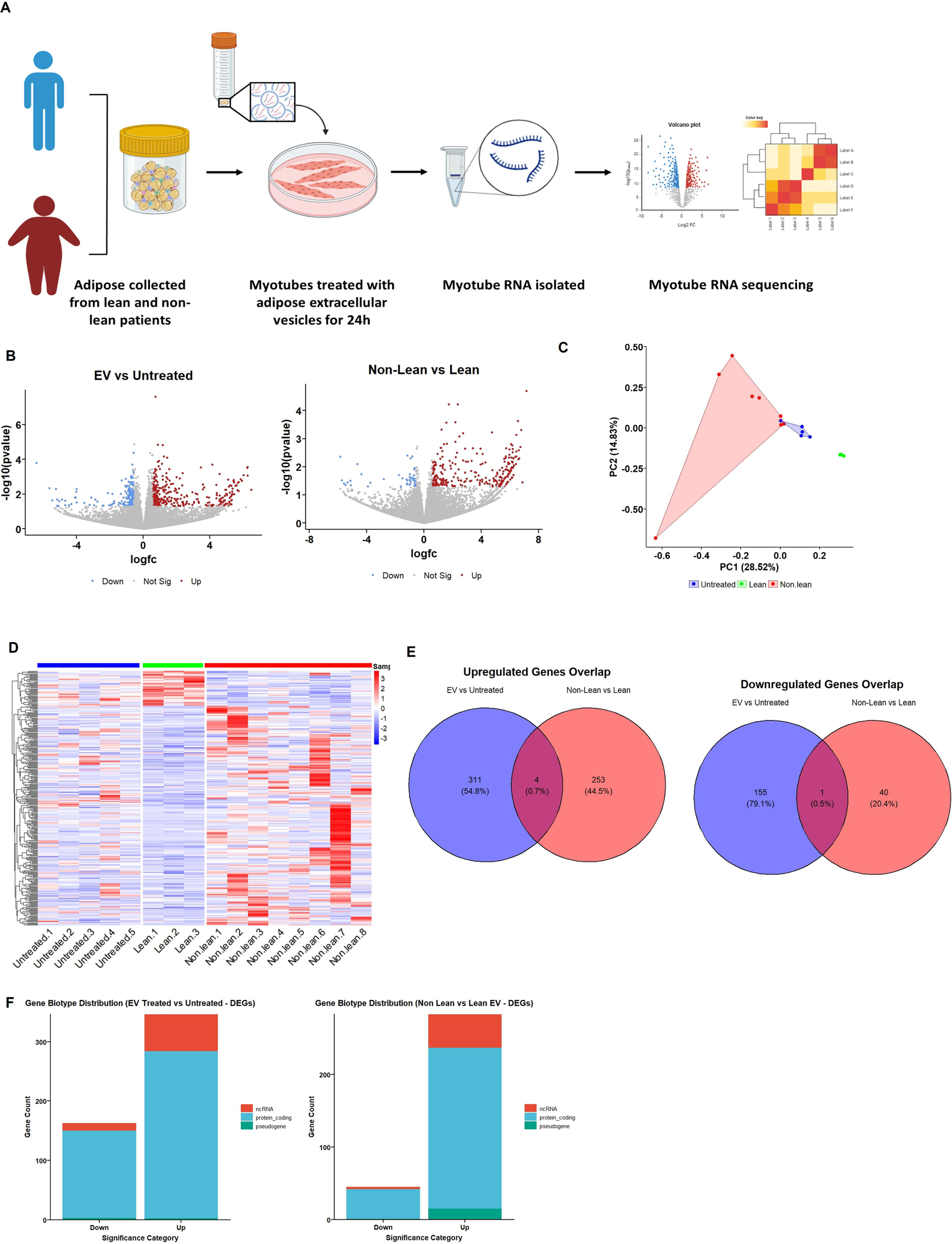
ACM-derived extracellular vesicles (EVs) modulate the transcriptome of human myotubes in a BMI-dependent manner. **(A)** Schematic of the experimental workflow. Differentiated human myotubes from a single donor were treated for 24 hours with EVs isolated from lean (BMI <25, n = 3) or non-lean (BMI >25, n = 8) adipose tissue, or left untreated (n = 5). These EVs were applied to independent myotube cultures, and RNA-seq libraries were generated from n = 5 untreated, n = 3 lean EV-treated, and n = 8 non-lean EV-treated myotube samples for downstream analysis. Total RNA was extracted and analysed by bulk RNA sequencing. **(B)** Volcano plots showing differentially expressed genes (DEGs) for EV-treated vs untreated myotubes (left) and non-lean vs lean EV-treated myotubes (right). Red: upregulated; blue: downregulated; grey: non-significant. **(C)** Principal component analysis (PCA) reveals clustering by treatment condition, with clear separation of non-lean EV-treated myotubes. **(D)** Heatmap of DEGs shows consistent expression profiles within each treatment group, highlighting the transcriptional effects of non-lean EVs. **(E)** Venn diagrams illustrate the overlap of significantly upregulated (left) and downregulated (right) DEGs between the two comparisons. **(F)** Gene biotype distribution of DEGs reveals that most upregulated genes are protein-coding, while non-coding RNA species (including lincRNAs and pseudogenes) are more prominent among the downregulated genes.

Minimal overlaps were observed between upregulated and downregulated DEGs of non-lean EV-treated and lean EV-treated myotubes compared to untreated, suggesting that the EV-induced transcriptional response in myotubes is both distinct from basal media exposure and highly dependent on the physiological origin of the EVs (lean vs. non-lean adipose tissue), with only 4 genes commonly upregulated and 1 gene commonly downregulated across both comparisons (Figure 3E).

To better understand the nature of differentially expressed transcripts, gene biotype distribution was examined. In both comparisons, the majority of DEGs were classified as protein-coding, with a smaller subset of non-coding RNAs (ncRNAs) also showing differential regulation (Figure 3F).

Gene ontology (GO) enrichment analysis provided further insight into the biological processes impacted by EV treatment. In the comparison of EV-treated vs. untreated myotubes, DEGs were enriched in pathways related to lipid metabolism and oxidative stress responses including “response to fatty acid,” and “regulation of reactive oxygen species metabolic process” (Figure 4A). These pathways suggest a potential EV-mediated reprogramming of cellular metabolism and stress handling.

**Figure 4.**
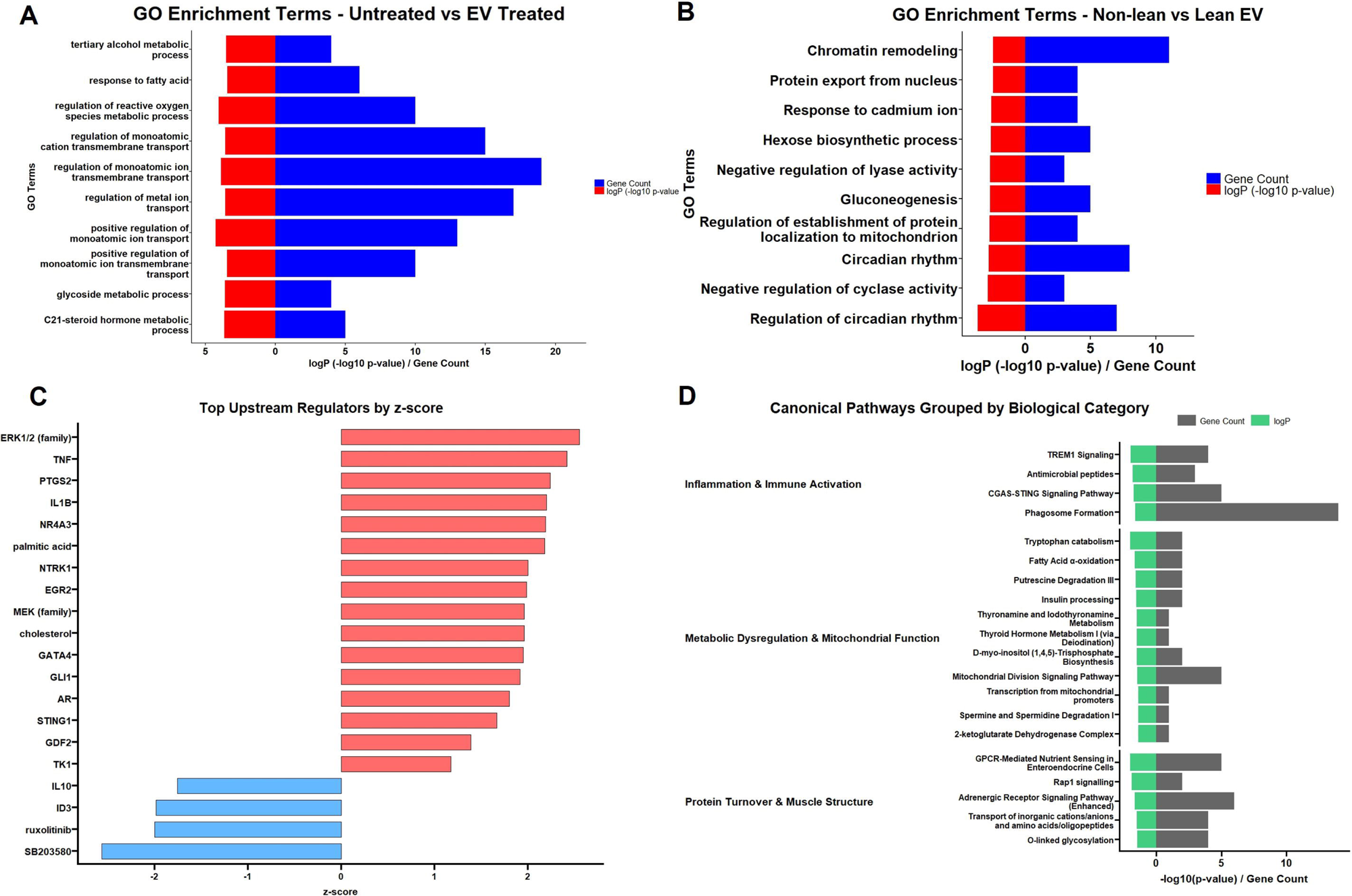
Gene ontology and pathway analysis of EV-driven transcriptional changes in human myotubes. **(A–B)** Gene ontology (GO) enrichment analysis of biological processes. DEGs from EV-treated vs untreated myotubes (**A**) are enriched in lipid metabolism, oxidative stress, and mitochondrial function pathways. DEGs from non-lean vs lean EV-treated myotubes (**B**) are enriched in chromatin remodelling, circadian rhythm regulation, and gluconeogenesis-related processes. **(C)** Predicted upstream regulators identified from the non-lean vs lean EV comparison. Pro-inflammatory cytokines such as TNF and IL1B are predicted to be activated, while compounds like SB203580 and ruxolitinib are predicted to be inhibited, suggesting potential targets for therapeutic modulation. **(D)** Canonical pathway enrichment grouped by functional category: (1) Inflammation and Immune Activation, (2) Metabolic Dysregulation and Mitochondrial Function, and (3) Protein Turnover and Muscle Structure. These pathways are enriched in DEGs from myotubes exposed to non-lean vs lean EVs, indicating broad transcriptional reprogramming of key muscle processes.

In contrast, the non-lean vs. lean EV comparison revealed enrichment in distinct biological processes such as “chromatin remodelling”, “protein export from nucleus”, “circadian rhythm”, and “gluconeogenesis” (Figure 4B). These data imply that non-lean EVs may alter nuclear and metabolic regulatory pathways in ways that lean EVs do not. Figure 4C shows predicted upstream regulators from transcriptomic differences between non-lean and lean adipose EV treatment of myotubes. This analysis identified several key inflammatory and stress-related regulators as activated in response to non-lean EVs, including TNF, IL1B, and PTGS2 (COX-2), as well as upstream signalling molecules such as ERK1/2 and NR4A3. These findings indicate that non-lean EVs elicit a transcriptional response consistent with heightened pro-inflammatory signalling, with TNF and IL1B being particularly important due to their central roles in muscle catabolism and insulin resistance.

The analysis also predicted that SB203580 (a p38 MAPK inhibitor) and ruxolitinib (a JAK inhibitor) would counteract the observed gene expression changes. This suggests that the pro-inflammatory effects of non-lean EVs could be reversed by inhibiting these pathways. Thus, SB203580 and ruxolitinib may represent potential therapeutic strategies to block EV-driven muscle atrophy.

Utilising IPA, we identified three core biological categories that undergo change following non-lean EV treatment compared to lean EV treatment. First, several inflammation and immune activation pathways are significantly enriched, including TREM1 signalling, CGAS-STING signalling, and phagosome formation. These findings suggest a heightened pro-inflammatory environment in response to non-lean EV treatment, consistent with the upstream regulator analysis shown in Figure 4C, where TNF and IL1B were among the top predicted activators. Second, pathways related to metabolic dysregulation and mitochondrial function are enriched, including fatty acid β-oxidation, thyroid hormone metabolism, mitochondrial division signalling, and tryptophan catabolism. These results indicate that non-lean EVs induce transcriptional changes associated with impaired metabolic flexibility and oxidative stress in skeletal muscle. Third, several pathways involved in protein turnover and muscle structure are highlighted, such as GPCR-mediated nutrient sensing, adrenergic receptor signalling, and protein glycosylation (Figure 4D).

To further explore the signalling consequences of EV treatment, a phospho-kinase array was performed on differentiated human myotubes following treatment with EVs from lean, and non-lean donors. This exploratory analysis revealed BMI-dependent phosphorylation changes in key signalling proteins, including eNOS, ERK1/2, EGF receptor, and STAT5a/b (Supplementary Figure 1E), consistent with the transcriptional reprogramming observed in response to EV exposure. Collectively, this suggests that non-lean EVs may disrupt structural integrity and protein synthesis processes in muscle cells, potentially contributing to muscle dysfunction.

Together, these findings highlight how differences in EV cargo from lean and non-lean adipose tissue impact key biological processes in skeletal muscle, encompassing inflammation, metabolism, and muscle structure.

### 3.4 Identification of adipose EV miRNA cargo

To identify regulatory mediators of the transcriptional and phenotypic changes observed in myotubes exposed to ACM-derived EVs, we profiled small RNA cargo from EVs isolated from lean (n = 3) and non-lean (n = 12) donors (Figure 5A). Small RNA sequencing identified 535 distinct miRNAs, of which 7 were differentially expressed between lean and non-lean EVs (log₂FC > 0.58, p < 0.05). Five miRNAs were uniquely detected in non-lean EVs, two were unique to lean EVs, and the remaining 528 were shared (Figure 5B–D). Among the differentially expressed miRNAs, miR-150-5p and miR-193b-5p were selected for downstream analysis due to their predicted involvement in muscle regulatory pathways and consistent upregulation in non-lean EVs [25].

**Figure 5.**
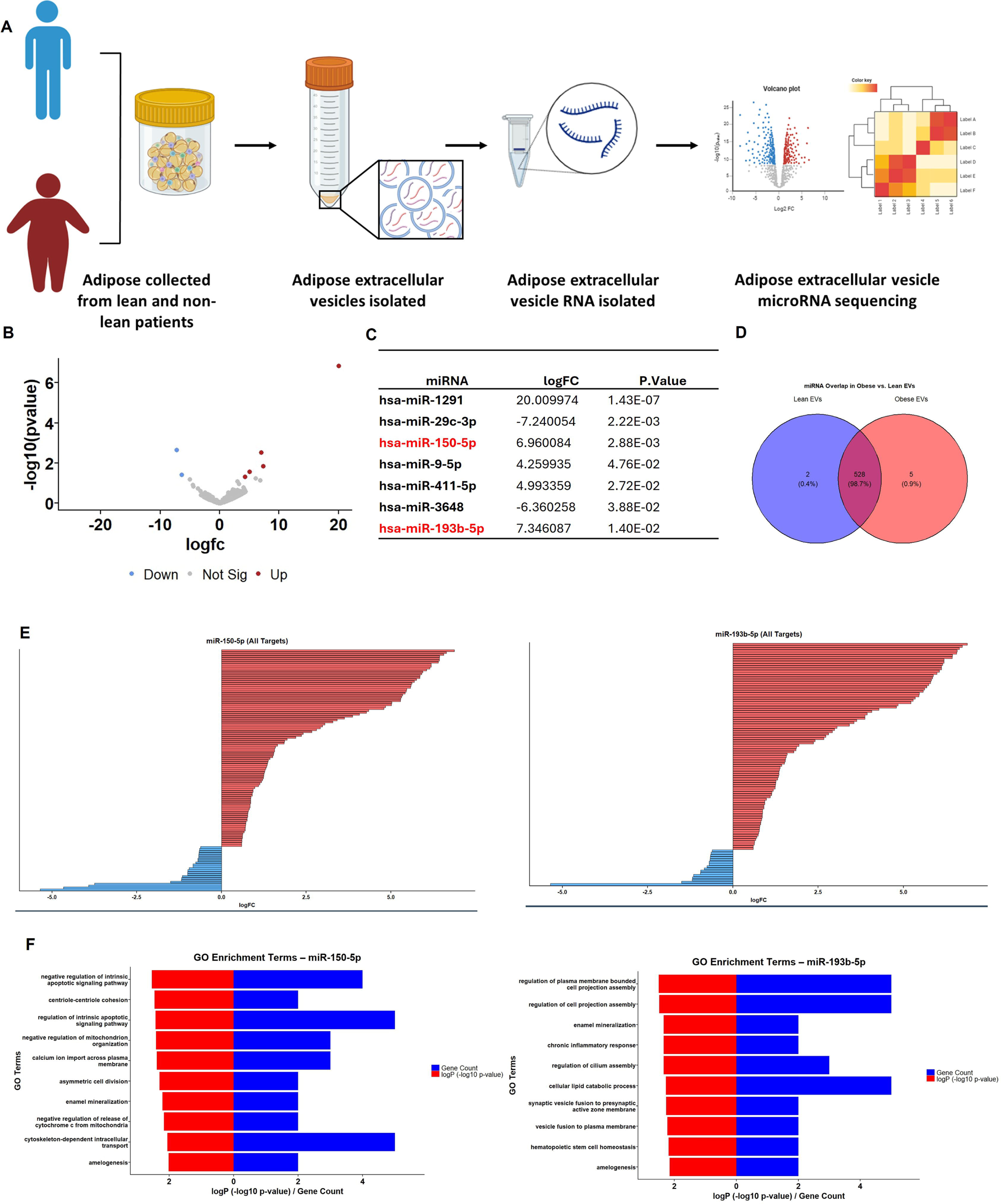
Small RNA sequencing of ACM-derived EVs identifies BMI-associated miRNAs with predicted gene targets linked to muscle atrophy pathways. **(A)** Schematic of the small RNA sequencing workflow. ACM-derived EVs were isolated from lean (BMI <25, n = 3) and non-lean (BMI >25, n = 12) donors. RNA was extracted and analysed by small RNA sequencing. **(B)** Volcano plot showing differentially expressed miRNAs between non-lean and lean EVs. Red: upregulated; blue: downregulated; grey: non-significant (log₂FC > 0.58, p < 0.05). **(C)** Table of significantly altered miRNAs between groups, highlighting both upregulated and downregulated candidates. miR-150-5p and miR-193b-5p were among the top upregulated miRNAs in non-lean EVs. **(D)** Venn diagram illustrating miRNA detection overlap. Most miRNAs (n = 528) were shared between lean and non-lean groups, with a small number unique to each condition. **(E)** Predicted mRNA targets of miR-150-5p (left) and miR-193b-5p (right) were cross-referenced with significantly differentially expressed genes (DEGs) from myotube transcriptomic data (non-lean vs lean EV treatment). DEGs consistent with miRNA-mediated repression (i.e., downregulated targets of upregulated miRNAs) are highlighted. **(F)** Gene ontology (GO) enrichment analysis of filtered miR-150-5p (left) and miR-193b-5p (right) target genes shows enrichment in biological processes including cell signalling, chromatin modification, inflammation, and homeostasis regulation.

To explore their potential effects on recipient cells, we used miRbase to collate the predicted target genes of miR-150-5p and miR-193b-5p and overlapped these with the transcriptomic data from non-lean EV-treated myotubes. Visualisation of fold-change values revealed broad downregulation among predicted miRNA gene targets, consistent with miRNA-mediated repression (Figure 5E). GO enrichment analysis indicated that miR-150-5p targets were associated with transcriptional regulation, cell proliferation, and glucose homeostasis, while miR-193b-5p targets were enriched for pathways related to ER stress, protein localization, and apoptosis (Figure 5F). These findings suggest distinct functional roles for each miRNA in shaping the cellular response to EV exposure.

Validation by qPCR in an independent cohort (n = 7 lean, n = 7 non-lean) confirmed the elevated abundance of both miR-150-5p and miR-193b-5p in non-lean EVs (p < 0.05). miR-155-5p, previously linked to muscle atrophy in obesity in murine models [26], was not differentially expressed in our sequencing dataset and showed no significant difference in validation (Figure 6B). miR-193b-5p has been shown to impair muscle growth in murine models of type 2 diabetes via inhibition of the PDK1/Akt signalling pathway [25]. Together, these results support the hypothesis that EV-associated miRNAs are selectively packaged in an adiposity-dependent manner and may contribute to the reprogramming of gene expression in skeletal muscle.

**Figure 6.**
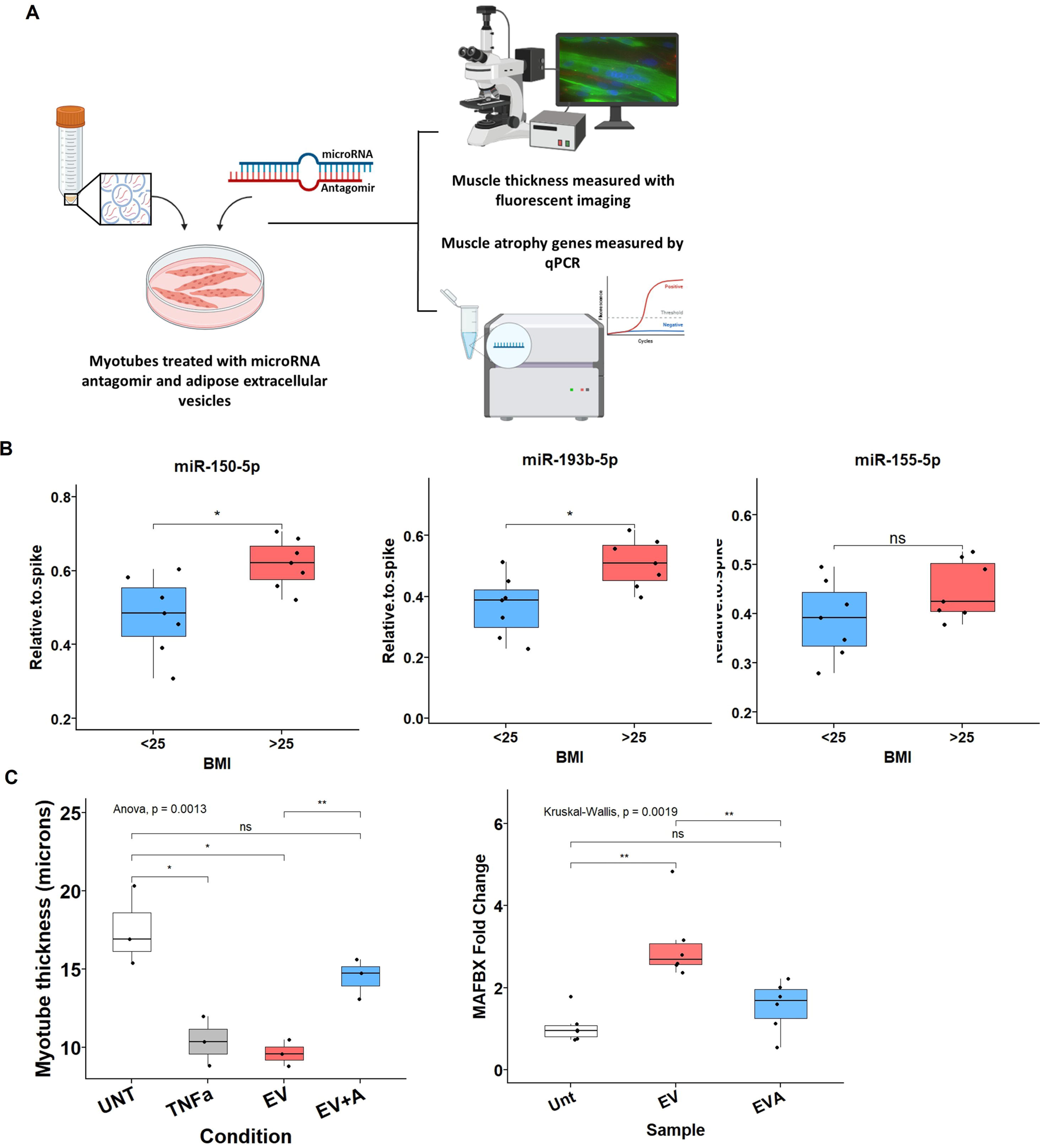
Inhibition of miR-150-5p attenuates EV-induced muscle atrophy in human myotubes. **(A)** Schematic of the experimental design. Differentiated human myotubes were treated with non-lean ACM-derived extracellular vesicles (EVs) in the presence or absence of a miR-150-5p antagomir (0.5 μM, 3′ cholesterol-conjugated; MedChemExpress #HYRI00301A). After 24 hours, myotube thickness was measured by fluorescent imaging and atrophy-related gene expression analysed by qPCR. **(B)** miRNA qPCR validation of EV cargo in a separate cohort of lean (n = 7) and non-lean (n = 7) ACM-derived EVs. miR-150-5p and miR-193b-5p were significantly upregulated in non-lean EVs, while miR-155-5p showed no significant difference (unpaired t-tests; each point represents an individual EV donor). **(C)** Functional effects of miR-150-5p inhibition. Left: Myotube thickness is significantly reduced by non-lean EVs (EV), but rescued by co-treatment with the miR-150-5p antagomir (EV + A). Data represent n = 3 replicate wells per treatment group from a single donor, with each point reflecting the average of multiple images. Statistical comparison was performed using one-way ANOVA with Holm–Bonferroni corrected post-hoc t-tests. Right: MAFbx expression is significantly upregulated by EV treatment and normalised with miR-150-5p inhibition (n = 6 wells per group). Statistical analysis used Kruskal–Wallis test with Wilcoxon post-hoc comparisons. All boxplots show median and interquartile range (IQR); whiskers extend to 1.5×IQR, and individual replicate values are plotted.

Next, to test functional involvement, we focused on miR-150-5p due to its consistent upregulation, regulatory target profile, and previously uncharacterized role in EV-mediated muscle atrophy. Differentiated human myotubes were treated with non-lean EVs in the presence or absence of a miR-150-5p antagomir (0.5 μM; cholesterol-conjugated) (Figure 6A). Non-lean EVs significantly reduced myotube thickness compared to untreated controls (mean ± SD: 9.6 ± 0.8 µm vs. 17.6 ± 2.5 µm, *p* < 0.01), replicating the atrophic effect seen in older donor-derived muscle. Co-treatment with a miR-150-5p antagomir partially rescued myotube thickness (14.5 ± 1.3 µm), representing a significant increase compared to EV-only treated cells (*p* < 0.01) (Figure 6C, left). This rescue was accompanied by a significant reduction in MAFbx (atrogin-1) expression, which was otherwise upregulated by non-lean EVs (Figure 6C, right). These findings establish miR-150-5p as a functional EV cargo component that contributes to both gene expression changes and atrophic phenotype induction in human myotubes, highlighting its potential as a mechanistic link between donor adiposity and muscle dysfunction.

## 4. Discussion

This study identifies a novel intercellular mechanism by which adipose tissue from non-lean (obese/overweight) individuals contributes to muscle atrophy. We also provide a detailed characterisation of EVs derived from primary human subcutaneous adipose tissue, including their size, concentration, and surface marker profile using multiple orthogonal platforms (NTA, ExoView, and CytoFLEX Nano). EVs isolated from non-lean ACM, induce human myotube atrophy and transcriptomic remodelling in an age-dependent manner. These effects, partially driven by miR-150-5p contextualise a mechanistic axis where adipose tissue communicates with skeletal muscle to drive atrophy and sarcopenia.

Functionally, non-lean EVs induced myotube atrophy and increased MAFbx expression (Figure 2), effects not observed with EV-depleted ACM. This highlights EVs as the primary mediators of atrophic signalling in adipose-muscle crosstalk. Notably, the atrophic phenotype was restricted to myotubes derived from older adults, suggesting age-related susceptibility to obese adipose EV signalling providing mechanisms that could underpin sarcopenic obesity. The use of primary human skeletal muscle cells enhances the physiological relevance of our model compared to traditional C2C12 systems, enabling direct investigation of human-specific responses to adipose-derived EVs, a level of translational relevance absent from murine and immortalised models [27].

Transcriptomic profiling revealed broad changes in gene expression linked to proteolysis, oxidative stress, and inflammation (Figure 3). While both lean and non-lean ACM-derived EVs induced transcriptional responses, non-lean origin produced a distinct pro-inflammatory and catabolic profile. IPA grouped enriched pathways into three categories: inflammation and immune activation, metabolic dysregulation and mitochondrial impairment, and altered protein turnover. Canonical pathways included IL-6, TNF, TLR and JAK/STAT signalling, while TNF and IL1B were predicted as upstream regulators (Figure 4). These transcriptomic findings were strongly supported by phospho-kinase profiling, which confirmed activation of key pro-inflammatory signalling cascades, including ERK1/2, STAT5a/b, and GSK-3 in a BMI-dependent manner (Supplementary Figure 1E). The activation of inflammatory pathways is particularly noteworthy given the established role of chronic low-grade inflammation in driving muscle wasting, insulin resistance, and impaired regeneration in ageing and metabolic disease. Our findings mirror prior observations that EVs can engage ERK1/2 and STAT pathways in muscle [28], but extend this by demonstrating that such signalling can be EV-induced specifically from non-lean adipose sources. This reinforces the concept that obesity-associated EV signalling may act as a dynamic trigger for catabolic and inflammatory cascades in human muscle tissue. To our knowledge, this is the first study to demonstrate that EVs derived from primary human subcutaneous adipose tissue can directly induce a pro-inflammatory and atrophic transcriptional programme in primary human skeletal muscle myotubes. Prior studies have typically employed adipocyte cell lines; our approach more accurately reflects physiological interactions relevant to human muscle atrophy [20, 25].

To explore mechanistic drivers, we profiled EV miRNA cargo. miR-150-5p and miR-193b-5p were consistently upregulated in non-lean EVs (Figure 5), with findings validated by qPCR (Figure 6B). While miR-155-5p has been linked to inflammation in obesity in murine models [28], it was not differentially expressed in our data. We integrated predicted miRNA targets with differentially expressed genes from our RNA-seq dataset (Figure 5E). Based on expression pattern, predicted impact on inflammatory pathways, and novelty miR-150-5p was prioritised for inhibition studies. Inhibiting miR-150-5p during EV treatment partially reversed myotube atrophy and MAFbx upregulation (Figure 6C), supporting its functional contribution. However, the incomplete rescue implies that other miRNAs, such as miR-193b-5p which was also identified in our study, or non-miRNA cargo, including proteins or lipids, may also mediate EV-induced effects.

Several limitations should be acknowledged. We did not conduct gain-of-function experiments using miRNA mimics, due to known challenges with cholesterol-conjugated mimic uptake and poor transfection efficiency in primary human myotubes [29, 30].s Due to limitations on sample volume, our study employed a single 24-hour treatment time point, limiting insight into temporal dynamics. We also observed discrepancies in EV concentration estimates across NTA, ExoView, and CytoFLEX Nano platforms. This reflects methodological differences, underscoring the challenges of EV quantification. Finally, this study focused on miRNA cargo and therefore did not investigate other EV components such as protein and lipids which may yield interesting results in future studies.

Future studies should explore whether agents like Ruxolitinib or p38 MAPK inhibitors counteract EV-driven atrophic effects. Interventions such as GLP-1 receptor agonists or structured exercise may attenuate the detrimental EV impact on muscle. Finally, our *in vitro* model provides a platform for testing whether physiological stimuli can enhance muscle resilience to pathogenic EV signalling. One such approach is electrical pulse stimulation (EPS), a method that mimics contractile activity *in vitro* and has been used to model the effects of exercise on muscle cells.

In conclusion, this study demonstrates that EVs from non-lean adipose tissue communicate pathogenic signals to skeletal muscle through miRNAs, particularly miR-150-5p, promoting phenotypic and transcriptomic changes consistent with atrophy. These findings offer mechanistic insight into adipose-muscle crosstalk in sarcopenic obesity and identify novel targets to preserve muscle mass and function in obese and ageing populations.

## Acknowledgments

The authors would like to thank the Dubrowsky legacy foundation for funding and support for JP. The authors would like to thank MyAge for supporting parts of this project. This study has been delivered through the National Institute for Health and Care Research (NIHR) Birmingham Biomedical Research Centre (BRC). The views expressed are those of the author(s) and not necessarily those of the BRC, the NIHR, the Department of Health and Social Care or any of our other funders.

## Ethics Statement

This study was approved by the UK National Research Ethics Service (reference: 16/SS/0172). Human skeletal muscle and adipose tissue samples were obtained from older adults undergoing elective orthopaedic surgery, with written informed consent obtained from all participants or their legal representatives. All procedures involving human participants were conducted in accordance with the ethical standards of the institutional and national research committees and with the 1964 Helsinki declaration and its later amendments. This study was conducted in compliance with the ethical guidelines for authorship and publishing in the *Journal of Cachexia, Sarcopenia and Muscle*.

