## Supplementary for "Obesity reprograms adipose extracellular vesicles to induce muscle atrophy via miR-150-5p-mediated transcriptional silencing"

**Methods:**

**Isolation of primary human myoblasts and their differentiation into multinucleated myotubes**

Satellite cells were isolated from skeletal muscle tissue. Briefly, excess adipose and connective tissue was removed from muscle samples, before being thoroughly minced with a scalpel and digested in 5 mL 0.05% trypsin-EDTA (in PBS) for 15min at 37°C. 5 mL myoblast growth medium (Ham's F10 with 2mM L-glutamine, 100 µg/mL penicillin/streptomycin, 20% FBS) was added and the digest centrifuged at 500 x *g* for 5 min. The resulting pellet was resuspended in myoblast growth medium and incubated for 20 min at 37°C, 5% $CO^2$ in an uncoated T75 tissue culture flask to deplete co-isolated fibroblast populations via preferential adherence. The cell suspension was then transferred to flasks or plates coated with 0.2% gelatin solution (in PBS) and the myoblasts cultured in growth medium. Myoblasts at 80% confluence were differentiated into myotubes by progressive serum reduction within culture media. Initially, for 24 hours, cells were cultured in myoblast growth media reduced to 10% FBS, before transfer to differentiation medium (Ham's F10 with 2mM L-glutamine, 100 µg/mL penicillin/streptomycin, 6% horse serum) for 8 days. All cells used in experiments were from passages 3 or less.

**CytoFLEX nano**

Reference bead standards (Beckman Coulter nanoVIS sizing beads) were employed to calibrate light scatter data to diameter (nm) using the FCMPASS software and Mie-scattering modelling with known refractive indices (NIST-traceable standards ranging from 80 nm to 450 nm). The calibration derived a collection half-angle for accurate size estimation on the VSSC1 channel, as per the manufacturer's protocol. The low refractive index setting was selected in FCMPASS to model biological EVs accurately. This approach follows the MIFlowCyt-EV guidelines for robust and reproducible EV sizing [30]. Prior to analysis, EV samples were diluted 1:600 in DPBS. DPS was filtered through a 0.02 µm Anotop® 10 syringe filter (Whatman, Merck, #WHA68091002) to minimize background interference. For CMO labelling, 10 µl of Cell Mask Orange (55 µg/ml stock in DMSO, Thermo Fisher Scientific) was added to 100 µl of EV sample and incubated for 1 hour at room temperature. Subsequently, 1 ml of filtered DPBS was added. As a control, 10 µl of CMO was incubated with 100 µl of filtered DPBS under identical conditions. Post-incubation, labelled samples were processed through qEVsingle/35 nm Gen 2 columns (IZON) using the IZON Automated Fraction Collector (AFC), and the first four post-void volume fractions were collected. These were analysed undiluted on the CytoFLEX nano.

**Transmission Electron Microscopy**

For transmission electron microscopy (TEM), carbon-coated 300 mesh nickel grids (TAAB Laboratories) were glow-discharged for 30 sec at 15 mA using an Agar Turbo Carbon Coater. 8µl of EV suspension was applied to each grid and incubated for 2 minutes at room temperature before excess was removed with filter paper. Grids were then immediately stained with 10 µl of 2% w/v uranyl acetate (Ladd Research) and blotted. After air-drying, grids were imaged using a Jeol JEM-1400Flash microscope with a Gatan OneView 16-megapixel camera at 100 kV.

**Proteome Profiler Phospho-Kinase Array**

Phospho-kinase profiling was performed using the Human Phospho-Kinase Array Kit (R&D Systems) according to the manufacturer's instructions. Differentiated human myotubes were treated for 24 hours with ACM-derived EVs isolated from donors categorized as lean (BMI <25), overweight (BMI 25–29.9), or obese (BMI ≥30). Cell lysates were prepared using the kit's lysis buffer, and equal amounts of protein were applied to the array membranes. After incubation with detection antibodies and chemiluminescent reagents, signal intensities were visualized using a ChemiDoc Imaging System (Bio-Rad). Spot intensities were quantified using ImageJ software, with background subtraction. Fold changes in phosphorylation levels were calculated relative to the untreated control group.

**PCR**

For confirmation of miRNA presence RNA was isolated from adipose EV. The TaqMan® Advanced miRNA Assay was used to confirm miRNA presence and was used as previously described (REF). Briefly, cel-miR-39-3p RNA spike-in (Qiagen, 339390) was added to each miRNA sample, the 3' end of all miRNA was then non-specifically polyadenylated. A universal adaptor was ligated to the 5' of all polyadenylated miRNA and was subsequently reverse transcribed using an oligo-dT primer 50-extended with a universal adapter sequence, that allows the addition of this adapter sequence in the final reverse transcription product (at the 3' end). Mir-Amp was then used to amplify the miRNA by using the 5' and 3' universal adapter sequences as hybridisation sites. The resultant concentrated cDNA template was then diluted 1:10 in RNase free water. Diluted cDNA miRNA templates underwent qPCR with compatible 2x Fast Advanced Master Mix and of 20X TaqMan® Advanced miRNA assays for miR-150-5p, miR-193b-5p, miR-155-5p and cel-miR-39-3p. Data are expressed relative to cel-miR-39-3p expression. mRNA levels of MAFbx and MuRF1 were determined by RT-qPCR, relative to the housekeeping gene 18S, using the iTaq™ Universal One-step kit mastermix (Bio-Rad). 5ng of RNA in a total reaction volume of 5 µl was used and all RT-qPCRs were performed in triplicate. Data was acquired using a Bio-Rad CFX Opus (Bio-Rad) and analysed by the 2-ΔΔCt method.

Supplementary Figure 1. Characterisation of extracellular vesicles (EVs) from adipose-conditioned media (ACM) and downstream phospho-proteome profiling.

**(A)** FCMPASS calibration report for CytoFLEX Nano (Beckman Coulter), generated using nanoVIS reference beads (Beckman Coulter) and low refractive index modelling to derive collection angle and calibrate VSSC1 light scatter to size (nm). Calibration supports accurate size and concentration estimations of EVs.

**(B)** Representative transmission electron microscopy (TEM) images of EVs from lean and non-lean ACM, negatively stained with uranyl acetate. Images show typical EV morphology, including spherical structures with lipid bilayer membranes. Comparable vesicle appearance between lean and non-lean samples is evident.

**(C)** Representative ExoView images showing EV capture spots for CD63-AF647 (red), CD9-AF488 (blue), CD81-AF555 (green), and Mouse IgG isotype control (no staining), confirming tetraspanin marker expression and effective capture of EVs from ACM. Comparable capture and marker intensity observed between lean and non-lean samples.

**(D)** Representative nanoscale flow cytometry plots (CytoFLEX Nano) illustrating sample purity after SEC cleanup. Left: CMO dye only control; Right: CMO-labelled EVs post SEC using Izon AFC and qEVsingle 35 nm columns. CMO+EV sample shows clear signal above dye-only background, validating labelling specificity and SEC cleanup effectiveness.

**(E)** Phospho-proteome profiling of differentiated myotubes treated with ACM-derived EVs. Fold-change in phosphorylation levels of selected signalling proteins (e.g., eNOS, ERK1/2, STAT5ab) compared to untreated control (UNT). Treatments included EVs from normal weight (NW), overweight (OW), and obese (OB) donors. Data indicate BMI-dependent modulation of signalling pathways associated with inflammation, proliferation, and cytoskeletal dynamics. Note: Proteome profiler data derived from one representative array per condition (n=1), presented for qualitative illustration.

S1:

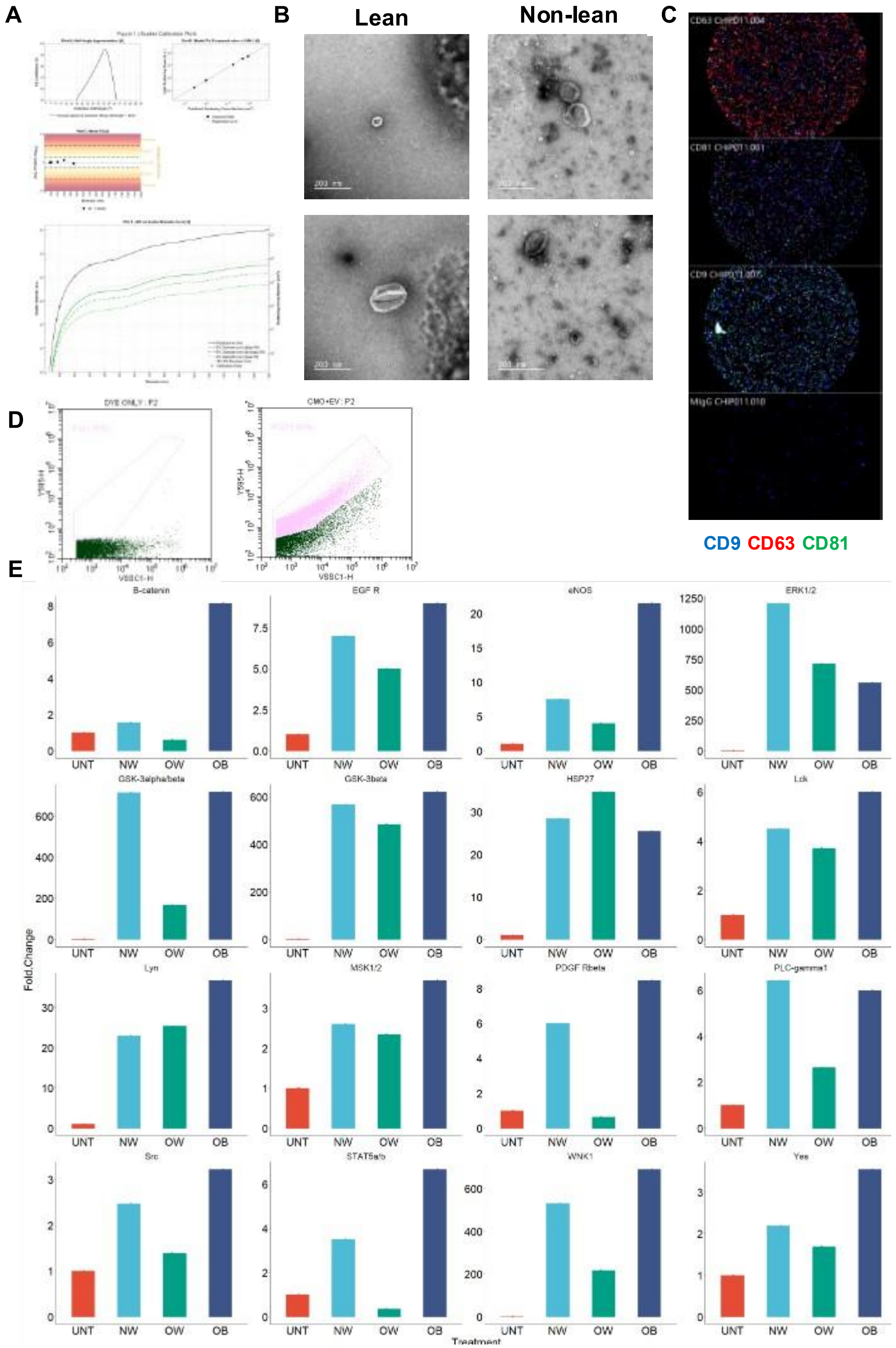

A
B
Lean
Non-lean
C
CD9 CD63 CD81
D
DYE ONLY : P2
CMO+EV : P2
VSSC1-H
Y585-H
E
B-catenin
EGF R
eNOS
ERK1/2
GSK-3alpha/beta
GSK-3beta
HSP27
Lck
Lyn
MSK1/2
PDGF Rbeta
PLC-gamma1
Src
STAT5a/b
WNK1
Yes
UNT
NW
OW
OB
Fold.Change
Treatment
